# STEM: A Method for Mapping Single-cell and Spatial Transcriptomics Data with Transfer Learning

**DOI:** 10.1101/2022.09.23.509186

**Authors:** Minsheng Hao, Erpai Luo, Yixin Chen, Yanhong Wu, Chen Li, Sijie Chen, Haoxiang Gao, Haiyang Bian, Lei Wei, Xuegong Zhang

## Abstract

Profiling spatial variations of cellular composition and transcriptomic characteristics is important for understanding the physiology and pathology of tissues in health or diseases. Spatial transcriptomics (ST) data are powerful for depicting spatial gene expression but the currently dominating high-throughput technology is yet not at single-cell resolution. On the other hand, single-cell RNA-sequencing (SC) data provide high-throughput transcriptomic information at the single-cell level but lack spatial information. Integrating these two types of data would be ideal for revealing transcriptomic landscapes at single-cell resolution. We developed the method STEM (SpaTially aware EMbedding) for this purpose. It uses deep transfer learning to encode both ST and SC data into a unified spatially aware embedding space, and then uses the embeddings to infer the SC-ST mapping and predict pseudo-spatial adjacency between cells in the SC data. Semi-simulation and real data experiments verified that the embeddings preserved the spatial information and eliminated technical biases between SC and ST data. Besides, we can use attribution analysis in STEM to reveal genes whose expressions dominate spatial information. We applied STEM to data of human squamous cell carcinoma and of hepatic lobule to uncover the spatial localization of rare cell types data and reveal cell-type-specific gene expression variation along a spatial axis. STEM is a powerful tool for mapping SC and ST data to build single-cell level spatial transcriptomic landscapes, and can provide mechanistic insights into the spatial heterogeneity and microenvironments of tissues.

## Introduction

High-resolution single-cell gene expression with spatial information is critical for revealing the mechanisms of cellular organization, embryogenesis, and tumorigenesis^1–5^, and could further enable therapeutic developments^6,7^. Recently, spatial transcriptomic (ST) profiling protocols have been rapidly developed and applied to study gene expression in spatial contests of many tissues^8–11^. The most commonly-used ST protocol aggregates multiple cells into one spot, providing in-situ gene expressions and spatial coordinates in a limited resolution^12^. On the contrary, single-cell RNA-sequencing (SC) data provide high-throughput gene expression profiles at single-cell resolution. They have advantages in inferring cellular identity, cell states, and trajectories of diverse cell types^13–16^, but lack spatial information.

It is desirable to computationally integrate SC and ST data to retain the advantages from both sides to facilitate comprehensive studies of spatial heterogeneities and variations of transcriptomics^17^. The current widely applied integration method is deconvolution, which employs SC or bulk RNA-seq data as references to estimate the cell type proportion of spots in ST data^18–21^. Referenced cell types should be specified before deconvolution, which limits the potential to discover sub-clusters or continuous variations within a cell type. Also, ST data are relatively less available compared with SC data. Deconvolution can only transfer the cell-type information in SC data to ST data, but cannot transfer the spatial information to SC data. This makes it hard to build the spatial single-cell transcription landscape based on the massive SC data. New methods are needed to transfer the spatial information in ST data to SC data. Such a single-cell level spatial landscape would greatly help investigate the spatial proximity of different cell types and spatial variations of gene expression for the same cell type.

The core of building a single-cell level spatial landscape is to establish SC-ST and SC-SC spatial associations. This task goes beyond merely mapping single cells onto ST data with high gene expression correlation, as several factors must be considered. Firstly, it is essential to align the data between SC and ST datasets. A good approach should address domain gaps such as batch effects and technical biases to ensure the accuracy of the results. Secondly, we need to filter the information on gene expression data to extract the part that reflects the spatial relationship between cells and spots. Gene expression profiles contain rich information on cell identity and state. We need to capture the information that is associated with spatial adjacency. Lastly, interpretability is critical to uncover the mechanisms governing tissue spatial organization. We need to identify the genes that determine the spatial location of individual cells.

To overcome these challenges, we propose STEM, a deep transfer learning model that learns SpaTially-aware EMbeddings of both SC and ST data for SC-ST and SC-SC spatial association inference. STEM features a shared encoder for SC and ST data to obtain their unified embeddings in the same latent space, and two predictors that simultaneously optimize these embeddings during the training stage. By preserving spatial information and eliminating domain gaps between SC and ST data, the optimized embeddings can be used to infer the SC-ST mapping and the pseudo-SC spatial adjacency. In experiments on both semi-simulation and real data applications, STEM outperforms existing methods in inferring spatial associations and preserving spatial topologies. We identified genes that dominate the spatial distribution of cells by interpreting the trained STEM model with the attribution technique. We used STEM to locate and reveal the spatial proximity of rare cell types in human squamous cell carcinoma (hSCC) data. We used STEM to construct the spatial transcriptomic landscape of hepatic lobules at the single-cell level and identified cell-type-specific gene expression variations along a spatial axis. STEM is a powerful method for revealing detailed and accurate maps of cellular spatial relationships, which can provide new mechanistic insights at the single-cell level into spatial transcriptomics studies.

## Results

### STEM: Learning spatially-aware embeddings of ST and SC data

Figure 1 illustrates the model of the STEM method. STEM has an encoder-predictor architecture and represents both ST and SC data as embeddings in a unified space. To address the issue of unstable and noisy representation of absolute spatial coordinates for cells that are too close or too far apart, we used a normalized spatial adjacency matrix between cells in SC data or spots in ST data as the prediction goal of STEM (Methods). In the training stage, we used the embedding of ST data and SC data to reconstruct two predicted spatial adjacency matrices. The ground truth spatial adjacency matrix is calculated according to the spatial coordinates of ST data. A cross-entropy loss is calculated on each pair of corresponding rows of the predicted and ground-truth matrixes.

**Figure 1.**
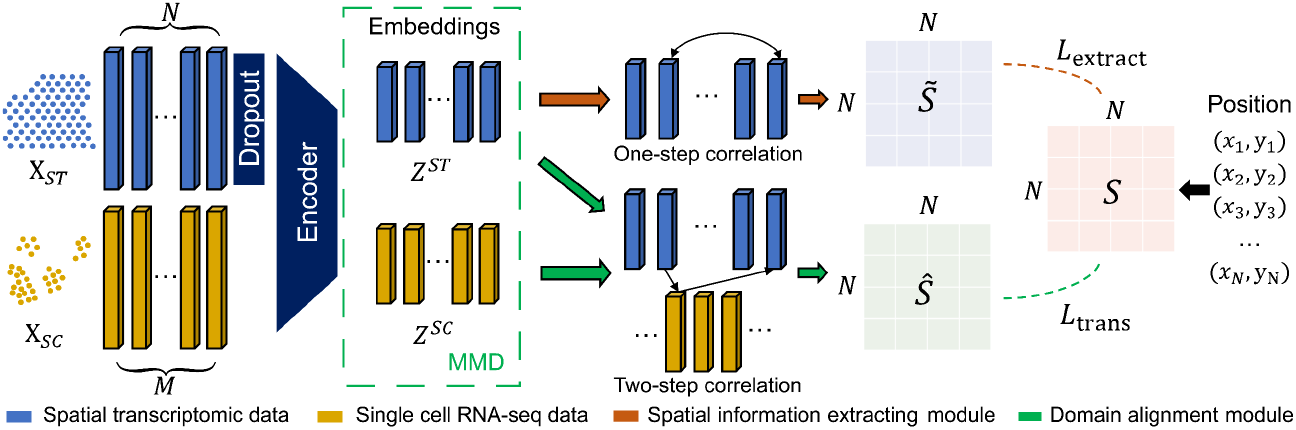
Schematic overview of STEM. Denoting SC and ST gene expression matrices as *X*_*ST*_ ∈ ℝ^*N*×*H*^ and *X*_*SC*_ ∈ ℝ^M× *H*^, where *N* and *M* are spot and cell numbers, and *H* is the number of genes. To align the sparsity of the *X*_*ST*_ with *X*_*SC*_, the *X*_*ST*_ first passes through an additional dropout layer. Then the processed ST matrix and the SC matrix pass through a shared encoder of STEM to get the corresponding unified embeddings Z^ST^ ∈ ℝ^h^ and *Z*^*SC*^ ∈ ℝ^*h*^ with the same hidden dimension size *h*, respectively. An MMD loss is used to align the distribution of SC and ST embeddings. These embeddings are used to predict the ST-ST spatial adjacency by two modules. The spatial information extracting module uses the correlation between *Z*^*ST*^ as the predicted ST-ST adjacency 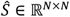. The domain alignment module uses the correlation between *Z*^*ST*^ and *Z*^*SC*^ to create the cross-domain mapping matrices which are multiplied to generate another ST-ST adjacency 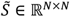. The reconstruction losses between the two predicted adjacency and the ground truth adjacency *LL*_*E*xtract_ and *LL*_trans_ are computed to optimize the STEM encoder and obtains the optimal embeddings. The ground truth spatial adjacency *S* ∈ ℝ^*N×N*^ is generated from the spatial coordinate of ST data.

Two predicted spatial adjacency matrices are obtained through two non-parameter predictor modules: the spatial-information extracting module and the domain alignment module. The spatial-information extracting module builds up the ST-ST predicted matrix by computing the correlations between the embeddings of spots in ST data. Each row of the matrix is normalized to 1, and thus each row represents the relative distance of one spot to others. The domain alignment module first eliminates the domain gap between SC and ST embeddings by minimizing the maximum mean discrepancy (MMD)^22^, and then constructs an SC-ST mapping matrix and an ST-SC mapping matrix. Both matrices are computed by the correlations between the embeddings of single cells in SC data and spots in ST data, and are normalized in a similar way as the ST-ST predicted matrix. The SC-ST mapping matrix describes the relative distance of one single cell to all spots, and the ST-SC is the reverse. STEM constructs an ST-SC-ST predicted spatial adjacency matrix by multiplying these two mapping matrices.

Through minimizing the loss during the training procedure, both modules simultaneously optimize encoder parameters to achieve meaningful embeddings of SC and ST data. The spatial-information extracting module encourages the ST embeddings to only contain spatial information, while the domain alignment module encourages the SC embeddings to be similar to ST embeddings and contain reasonable spatial information for building the optimal mapping matrices. Unlike unsupervised dimension reduction algorithms such as autoencoder^23^ and VAE^24,25^ which condense all information into the latent space, STEM uses spatial adjacency to supervise the embeddings, helping to extract the spatial information from gene expression and also eliminate the domain gap between SC and ST data.

After training, the optimized ST-SC and SC-ST mapping matrices are used to build the SC-ST spatial adjacency, and the correlation within SC embeddings builds SC-SC spatial adjacency. Additionally, STEM can link the predicted spatial adjacency weights to gene expression. This is because the weight in the predicted spatial adjacency is generated from the embeddings which are encoded from gene expression. By following the spatial adjacency-embedding-gene path, it is possible to identify the genes that significantly contribute to determining the spatial location of each cell. We employ the integrated gradient technique^26^ to achieve this. Detailed descriptions of the STEM model and algorithm are provided in the Methods section.

### Semi-simulation experiments showed that STEM achieves accurate spatial mapping at both cell and tissue levels on embryo atlas data

To benchmark the performance of STEM, we conducted a semi-simulation experiments based on the synthetic data generated from the Spatial Mouse Atlas^27^ dataset. It is a single-cell resolution spatial transcriptomics dataset. The dataset was produced by seqFISH^28^ and contained three distinct mouse embryo slides. On each embryo slide, the single-cell level gene expression profiles were provided and each cell had a spatial coordinate. We treated the gene expression data as pseudo-SC data without the spatial coordinates, and treated the spatial coordinates as the ground-truth to test the predictions of methods. We synthesized pseudo-ST data to simulate characteristics of the widely used 10X Visium data: they had a resolution lower than the single cell level and covered only partial cells in the tissue (Figure 2A). Specifically, we created a grid on the tissue slide and generated pseudo spots at the intersections to cover a portion of single cells in the tissue slide. The gene expression values of each spot were the gene expression summation of all covered cells. The true cell-type proportion of each ST spot was computed based on the cell type annotations of covered single cells.

**Figure 2.**
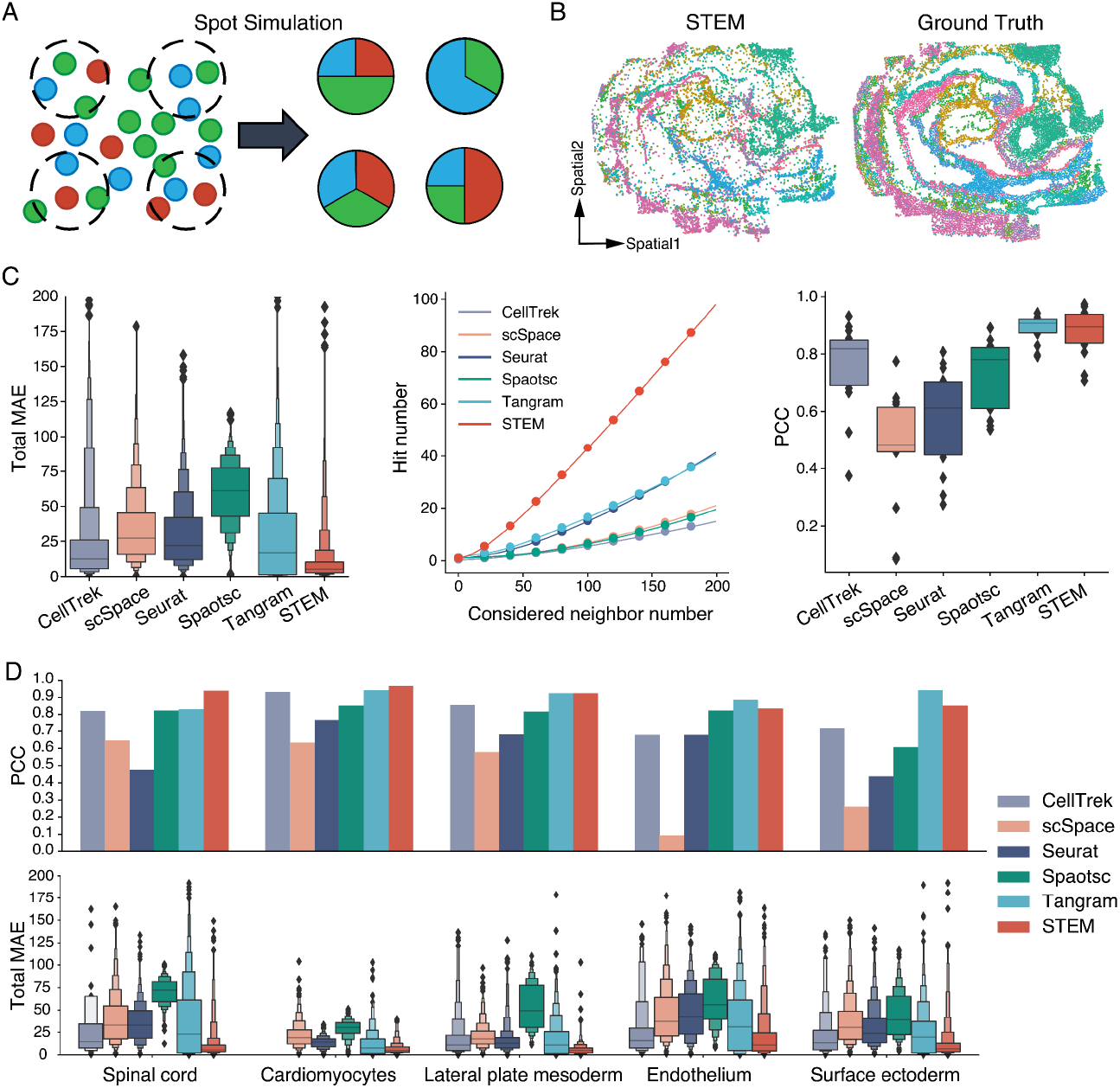
The performance evaluation results of different methods on a semi-simulation experiment using mouse embryos. A) An illustration of pseudo-ST data generation. Spots on ST data contain only a fraction of single cells. B) The reconstructed spatial distribution by STEM (Left) versus the ground truth spatial distribution (right). Colors indicate different cell types or regions. C) The mean absolute error (MAE), hit number and Pearson correlation coefficient (PCC) performance of different methods on the first mouse embryo data. The lower the MAE, the better. The higher the hit number and PCC, the better. D) The PCC and MSE performance of methods on five cell types. These manually selected cell types covered all comparison results (equal, lower and higher) between the PCC of Tangram and STEM. The bar plot shows the PCC between ground truth and predicted spatial distributions of five cell types (above). The enhanced boxplot shows the corresponding MAE of all methods (below). The enhanced boxplot has more quintiles and each edge of the box splits the rest data into two halves.

We applied STEM on the pseudo-SC and pseudo-ST data of each embryo slide to get the unified embeddings and construct the ST-SC mapping and SC-SC adjacency matrices. We compared STEM with the other five single-cell mapping methods: CellTrek^29^, scSpace^30^, Seurat^31^, Spaotsc^32^ and Tangram^33^. CellTrek and Seurat use mutual nearest neighbors to transfer the spatial coordinates from ST to SC. scSpace uses an MLP to predict the absolute spatial coordinates by gene expression data. Spaotsc uses the optimal transport theory to learn the SC-ST mapping matrix. Tangram directly learns a mapping matrix to convert SC to ST, and the matrix is optimized by minimizing the cosine similarity between the converted and ground truth ST. We uniformed the outputs of all methods into predicted spatial coordinates, SC-SC adjacency and SC-ST mapping for further evaluation (Methods).

We first evaluated whether the absolute spatial distribution of single cells can be reconstructed. The reconstruction results of all methods on three embryos were shown in Fig. 2B and Fig. S1-3. From these results, we can see that STEM is the only method that preserves the original topology structure of all single cells. CellTrek gave a similar shape, but it predicted spatial information for only about 38% of single cells, with the rest discarded by their algorithm. We used the mean absolute error (MAE) between the predicted and ground-truth spatial coordinate to measure the accuracy of predicted coordinates. CellTrek, scSpace, Seurat and Tangram achieved similar performances while Spaotsc had the highest error (Fig. 2C left and Fig. S4). STEM consistently achieved the lowest MAE compared to all the other methods.

We then validated the correctness of predicted SC-SC adjacency by computing the hit number, which is defined as the number of one cell’s true k-nearest neighbors that are successfully predicted. STEM got the highest hit number, about two-fold of that of the second best method on all three embryos. (Fig. 2C middle and Fig. S5) When considering 200 true neighbor cells, STEM identified approximately 100 correct neighbors on embryo 1 data. It is interesting to notice that while for most methods including STEM, a lower MAE corresponds to a higher hit number, CellTrek achieved the low MAE but got the lowest hit number. This might be caused by the point repulsion process used in their method.

We also evaluated whether the true cell-type spatial distributions can be correctly mapped from single cells to spots. We used the ST-SC mapping matrix to transfer the cell-type annotation from single cells to ST spots and thus each ST spot had a predicted cell proportion (Method). The Pearson correlation coefficient (PCC) was computed between the predicted and true distribution of cell types (Methods). STEM and Tangram had comparable average PCCs for all cell types across all three embryos (0.85 ± 0.02 and 0.87 ± 0.01, mean ± s.d.) (Fig. 2C right and S6). All other methods gave mean PCCs lower than 0.8. We found that the performance is also influenced by the properties of cell types (Fig. S7 and S8). For example, cardiomyocytes had obvious aggregation patterns on slides, making it easier to map their spatial distribution by just identifying the cell type from gene expression. As a result, all methods yielded good results on cardiomyocytes. Conversely, the distributions of hematoendothelial progenitors were scattered in space, necessitating a more detailed subdivision of cell subtypes for mapping cells into their locations. All methods achieved lower PCCs on this cell type.

We observed that Tangram and Spaotsc, which rely on the similarity of whole gene expressions, can recover the cell type spatial distributions but hardly estimate the cell coordinates well. On the five cell types shown in Fig.2D, Spaotsc reached the average PCC value of all methods, and Tangram had both high and low PCC values compared with that of STEM. But with the MAE metric, Spaotsc performed worse than the average, while Tangram was also inferior to STEM. This showed that the expression-based reconstruction method can locate the cell type in the corresponding position but was not sensitive enough to accurately locate the single cell. And these results also demonstrated that the similarity of raw gene expression was easily dominated by other information such as cell type identity, while STEM achieve more accurate single-cell spatial reconstruction by explicitly embedding spatial information.

Overall, these semi-simulation experiments systematically showed that STEM can accurately reconstruct the spatial landscape of all single cells in SC data by transferring information from ST data that cannot reach single-cell resolution. STEM consistently achieved more accurate single-cell spatial adjacency estimation compared with other methods.

### STEM builds spatial informative embeddings and identifies spatial dominant genes

Interpretability is important for using machine learning-based prediction methods to study underlying mechanisms. We interpreted STEM by analyzing the latent embeddings and traced genes that contribute to the spatial information of cells. As an example, we experimented on single cells in the forebrain, tegmentum, midbrain, and hindbrain regions of embryo 1 data. These regions were reported to have spatially-driven transcriptional heterogeneity^27^. The latent embeddings of cells obtained from the STEM encoder showed a clear trend from the forebrain to the hindbrain in the uniform manifold approximation and projection (UMAP) plot, verifying the ability of STEM for extracting spatial heterogeneity from gene expression profiles. The spatial topology in UMAP even preserved the hollow structure of the forebrain in the original slide (Fig. 3A). Similar structures were not captured in the reference UMAP plot, which was given by the PCA embeddings obtained by the standard pipeline of SCANPY^34^. These results suggest the importance of extracting spatial information from gene expression for the spatial mapping of SC data.

**Figure 3.**
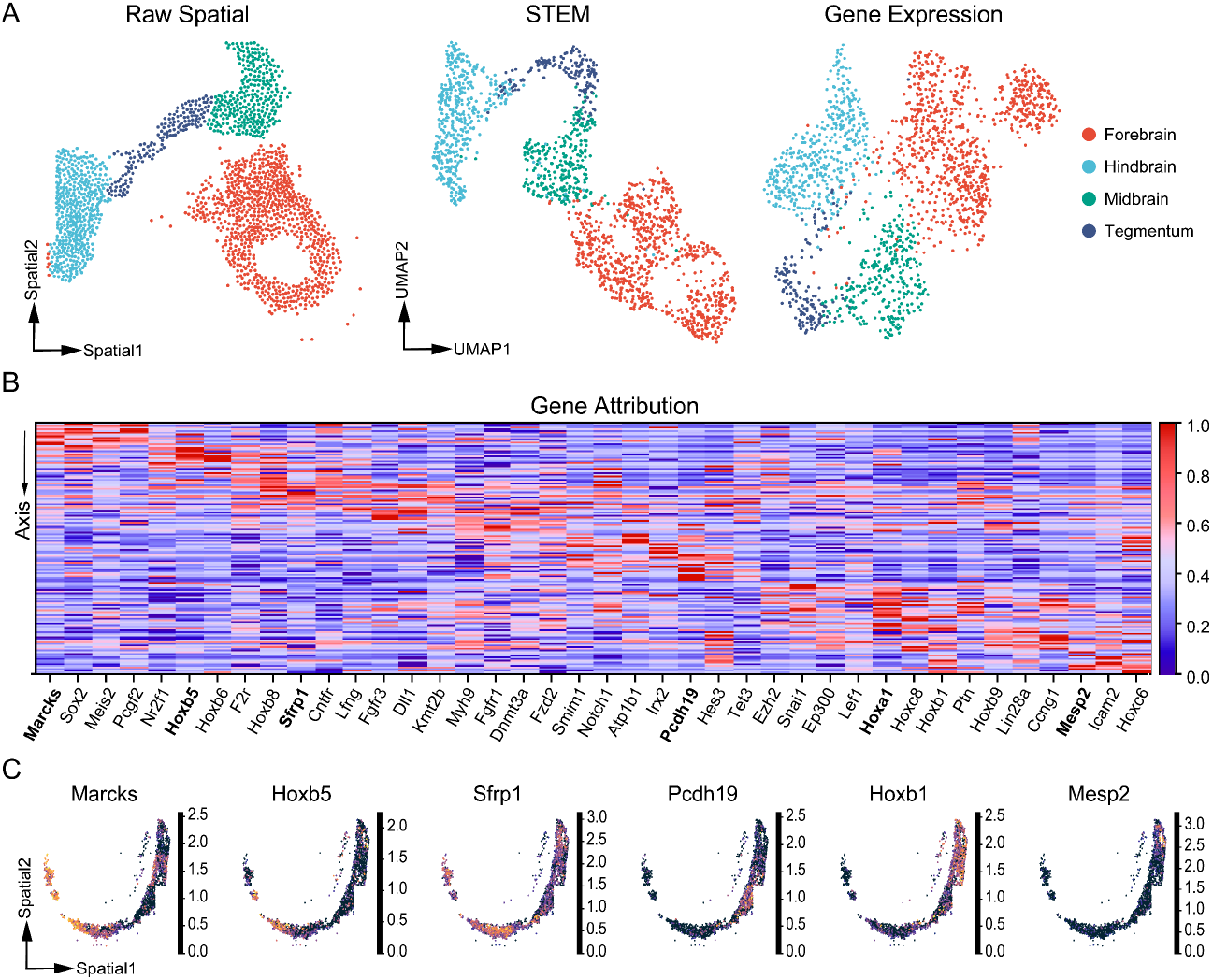
Interpretation of the STEM model. A) Raw spatial distribution of cells (Left) and a UMAP visualization of cell embeddings obtained from STEM and Gene expression. The color indicates the different spatial region annotation. B) The heatmap shows the attribution score of SDGs along the spinal axis. Each column represents a gene expression vector, with the attribution score scaled from 0-1. C) The spatial expression patterns of six SDGs are shown. From left to right, the genes are highly attributed in different regions, corresponding to the bolder name in the heatmap.

By utilizing integrated gradient techniques^26^ on the STEM model, we assigned each cell an attribution vector that showed the contribution of each gene in determining the spatial location of this cell (Methods). We focused on the spinal cord region which is spatially distributed along the anterior-posterior axis. We formulated a spatial trajectory on it and computed pseudo time score for each single cell. (Fig. S9). We divided the trajectory into 11 segments based on the score and identified spatial dominant genes (SDGs) using the Wilcoxon rank sum test. SDGs were defined as genes with significantly highly expressed attribution scores (FDR < 0.05) compared to other segments. A total of 272 SDGs were identified among 351 genes. The top differentially scored SDGs of all segments displayed a clear diagonal pattern in the heatmap (Fig. 3B), indicating that the attribution scores of these top SDG are highly expressed only in the specific spatial region. And these genes’ raw gene expression also exhibited similar expression patterns. For instance, along the axis, genes *Marcks, Hoxb5, Sfrp1, Pcdh19, Hoxb1* and *Mesp2* showed high gene expression values in the corresponding spatial segments with high attribution scores (Fig. 3C). There were also some SDGs that didn’t visually exhibit a strong spatial expression pattern, such as *Nebl, Bak1, Kmt2d, Suz12*, and *Fgfr2* (Fig. S10), which demonstrated that STEM can extract spatial information from genes that do not have easily identifiable spatial patterns.

Additionally, the identified SDGs had potential interests in revealing tissue organization mechanisms, as supported by previous studies. For instance, *Marcks* has been reported to be highly expressed in the nervous system and is important for the regulation of embryo development^35^. The Hox genes (*Hoxb5* and *Hoxb1* in our case) have been reported to emerge gradually from the posterior aspect of the vertebrate embryo and display anteroposterior positional information during tissue generation^36^. These results manifest that the attribution analysis on the STEM model made it possible to identify SDGs that have high contributions for determining cell location, which could provide insights into the spatial formation and evolution of cells in complex normal tissues or tumor microenvironments.

### STEM reconstructs single-cell spatial distribution on the human middle temporal gyrus

We applied STEM on the well-studied human middle temporal gyrus (MTG) region^37^ to verify the performance of STEM. The SC data we used were sequenced by the SMART-seq protocol and derived from 8 donors between 24 and 66 years old^38^. These SC data lacked the spatial coordinates of cells, but had the dissection information of brain subregions and manually annotated cell types. The ST data we used were at single-cell resolution produced by the *in-situ* sequencing technique MERFISH^39^. The ST data contained about 4,000 genes with spatial coordinates and layer segmentation information.

The single-cell spatial distribution reconstructed by STEM closely resembled the human cortex topological structure in the ST data (Fig. 4A), whereas other methods produced more blurry distributions (Fig. S11). We further examined the cortical-depth distribution of single cells estimated by all methods across different dissection layer groups. We divided single cells into 6 layer groups of L1-L6 based on the tissue dissection information. We divided the ST data into L1, L2/3, L4, L5 and L6 subregions using the reference provided in the original study. Ideally, cells from the same dissection layer group should aggregate in the corresponding layer spatial region on the MERFISH tissue. As depicted in Fig. 4B, the cortical-depth distribution of single cells produced by all methods exhibited laminar organization, meaning that cells in the L1 group had the minimal relative depth, while cells in the L6 group had the maximum relative depth. However, all other methods except STEM failed to locate the cells in their corresponding regions. Spaotsc and Tangram compressed all cells into L1 and L2/3 regions. Seurat and CellTrek located cells of L6 group in the shallow region, while scSpace located all cells with an offset to deeper region. Only the depth distribution estimated by STEM fitted well with the true layer regions identified from ST data.

**Figure 4.**
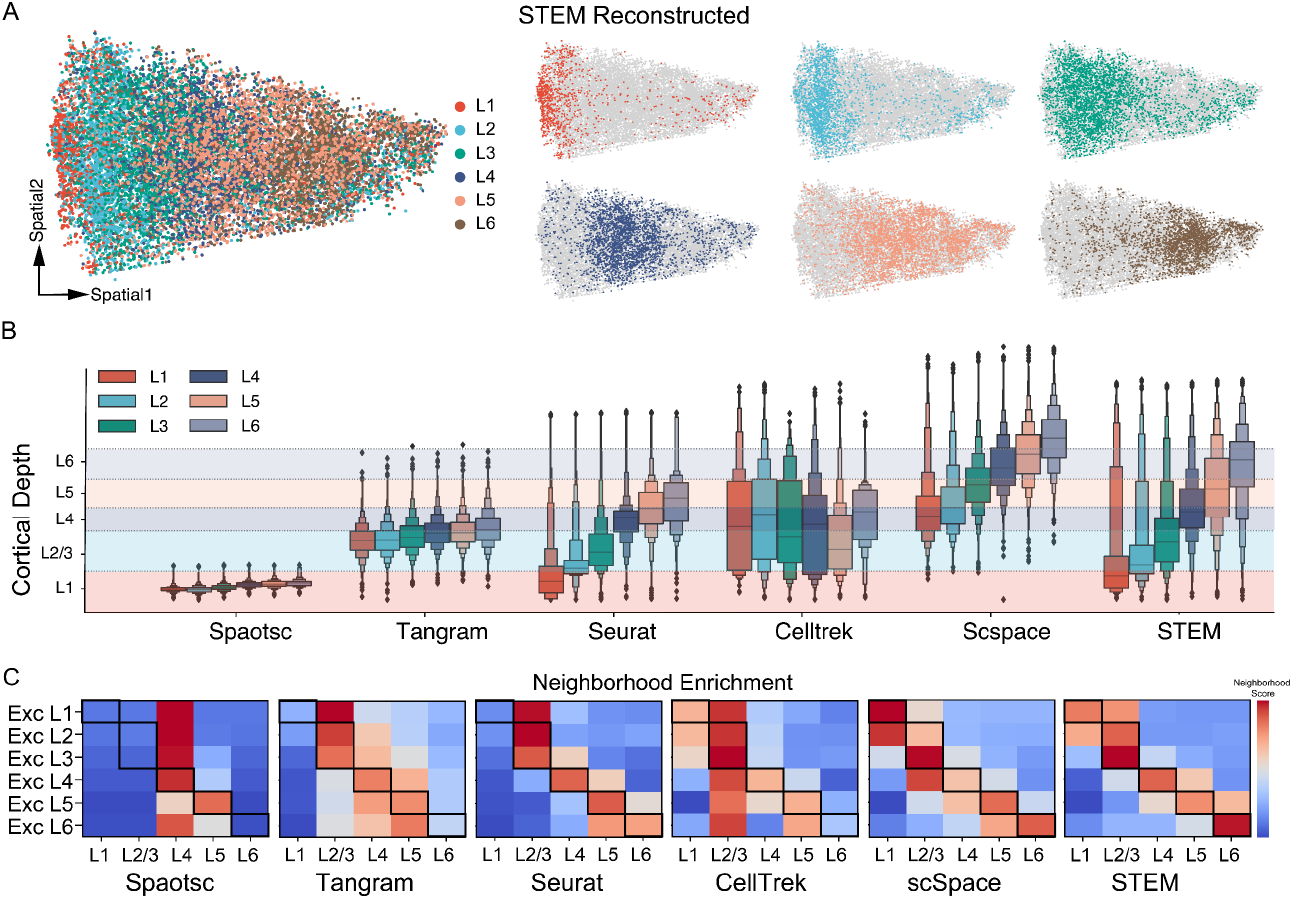
Performance evaluation on human MTG using all methods. A) The overall reconstructed spatial distribution of single cells obtained by STEM (left). The six subplots show the spatial distribution of cells in L1-L6 groups (right). These groups were determined based on tissue dissection information. B) Comparison of cortical-depth distribution of cell groups between different methods. The boxplot in various colors displays the cortical-depth distribution of different single cell groups. The dashed lines indicate the boundaries of layer regions given by ST data. C) Neighborhood enrichment analysis between SC and ST data using all methods. The x-axis represents regions in the ST data while the y-axis represents SC excitatory neurons from different dissection layers. The score is row-normalized, and the red color indicates a higher neighborhood score. Bold squares represent areas where high scores are expected.

We further validated this result by computing the neighborhood enrichment score^40^ between the SC and ST data. We focused on the excitatory neurons in the SC data. The results are shown in Fig. 4C. The STEM results showed a clear diagonal neighborhood score on the heatmap, indicating the estimated SC spatial distribution of all layers was in accordance with the ST ground-truth. Spaotsc mapped all cells around the L4 region. Tangram and Seurat failed to locate the L1 excitatory neurons which locate in a thin region in human MTG. scSpace mixed the L3 and L4 excitatory neurons. STEM was the only method that recovered the absolute spatial distribution and preserved the spatial topology.

### STEM locates tumor-specific keratinocytes and immune cells at the single-cell level

We applied STEM to real datasets to study tumor microenvironments (TME). The characterization of the spatial architecture and arrangement of single cells in the tumor microenvironment is critical for understanding tissue heterogeneity and plasticity^7,41,42^. We applied STEM to a human squamous cell carcinoma (hSCC) dataset^43^. We used the paired SC and ST data from the hSCC tissue of two donors. The SC data were obtained by 10X single-cell sequencing, and a detailed manual cell-type annotation was provided. The ST data were obtained by an early version of the 10X Visium (Spatial Transcriptomics) technique^12^ with HE-stained tissue images provided.

STEM mapped all the SC data to ST slides accompanied with hematoxylin and eosin (H&E)-stained histological images (Fig. 5A). We verified that the tumor specific keratinocytes (TSKs) were colocalized with endothelial cells at the top tumor leading edge in both donors (Fig. 5B). In the original study, the TSK localization was estimated by scoring each spot with TSK-signature genes which were manually identified from scRNA-seq data. Compared to this process, STEM achieved similar results but reduced the workload and the potential bias in the manual gene selection process. We further explored the spatial distribution of other keratinocyte (KC) subtypes (including tumor basal, tumor cycling, and tumor differentiating KCs). Neighborhood enrichment analysis (see Methods for details) showed that TSKs tended to spatially self-aggregate and separate from other KCs, especially far away from the tumor cycling KCs. And tumor differentiating cycling and tumor basal KCs were colocalized (Fig. 5C). This is consistent with the finding in the original study that TSKs and other KCs are distributed in different leading edges. These findings revealed the spatial characteristic of TSKs in the tumor microenvironment.

**Figure 5.**
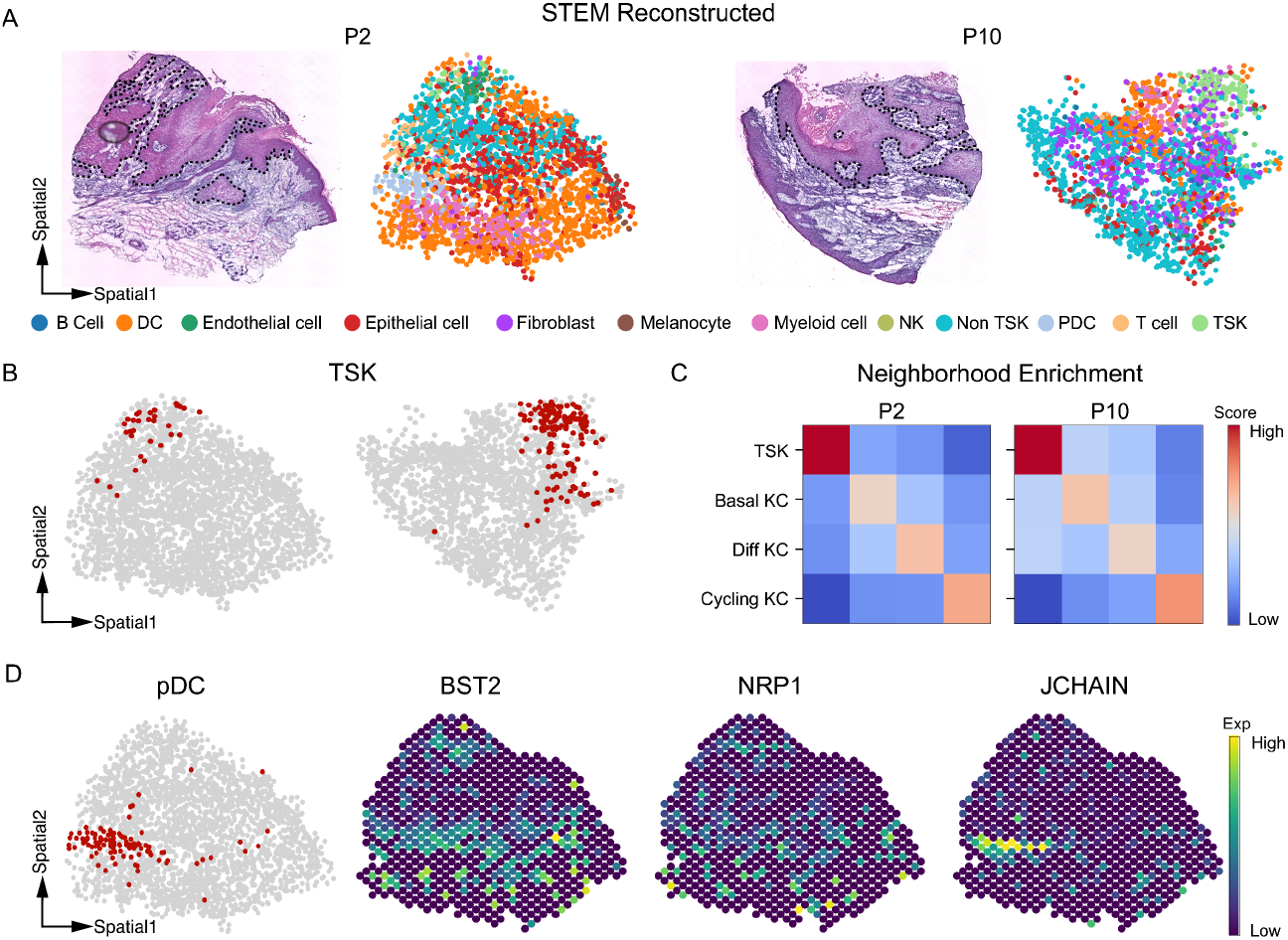
STEM results on human squamous cell carcinoma data. A) The HE images and spatial reconstruction results of STEM on patient 2 and patient 10. The black dashed line annotates the tumor-non tumor leading edge observed in the image. Colors of dots represent different cell types. B) Highlighted spatial distribution of TSK cells on P2 and P10 slides. C) Neighborhood enrichment analysis on tumor keratinocyte subtypes. The score is row normalized and thus asymmetric. Color in red indicates a higher neighborhood score. D) The spatial distribution of pDC cells and three spatial expression patterns of corresponding pDC and other immune-related marker genes.

We then studied the immune cell population in the non-TSK leading edge area, where we observed the predominant spatial positioning of plasmacytoid dendritic cells (pDCs) at the bottom leading edges in donor 2 (Fig. 5D). The expression levels of pDC and immune marker genes such as BST2, NRP1, and JCHAIN (obtained from CellMarker^44^) were also found to be significantly high in the same region (Fig. 5D). The original study had reported the activation of IFNs-related signaling pathways in this region. STEM allowed us to infer that the localization of pDCs could be the potential driving force of this pathway. This inference was consistent with previous studies that have shown that pDCs secrete significant amounts of type 1 interferon ^45^. Our results and analysis illustrate that STEM streamlines the determination of the spatial location of rare cell types without the need for manual identification of signature genes.

### STEM reveals cell-type transcriptomic variations along the liver zonation axis

We applied STEM to mouse livers data to characterize the cell-type-specific transcriptomic spatial variation in hepatic lobules. In the liver, hepatic lobule is a repeated basic anatomical unit ^46^ that displays a spatial trend from the portal vein (PV) to the central vein (CV). Studying the transcriptomic variation of different cell types along the trend is critical for revealing the mechanism of liver diseases such as cirrhosis and hepatocellular carcinoma^47^. We used scRNA-seq and ST data from a liver cell atlas^48^. Four liver tissue slide datasets of a healthy mouse from 10X Visium technique were used as the ST data. Single cells from all health mouse are used as the SC data. Each spot in the ST data was manually assigned a zonation score that reflects the scaled distance from the spot to the PV. Using the spatial mapping matrix obtained from STEM, we reordered the cells along the lobular axis from PV to CV, and revealed the spatial variation of cell-type-specific gene expression along this axis (Fig. 6B).

**Figure 6.**
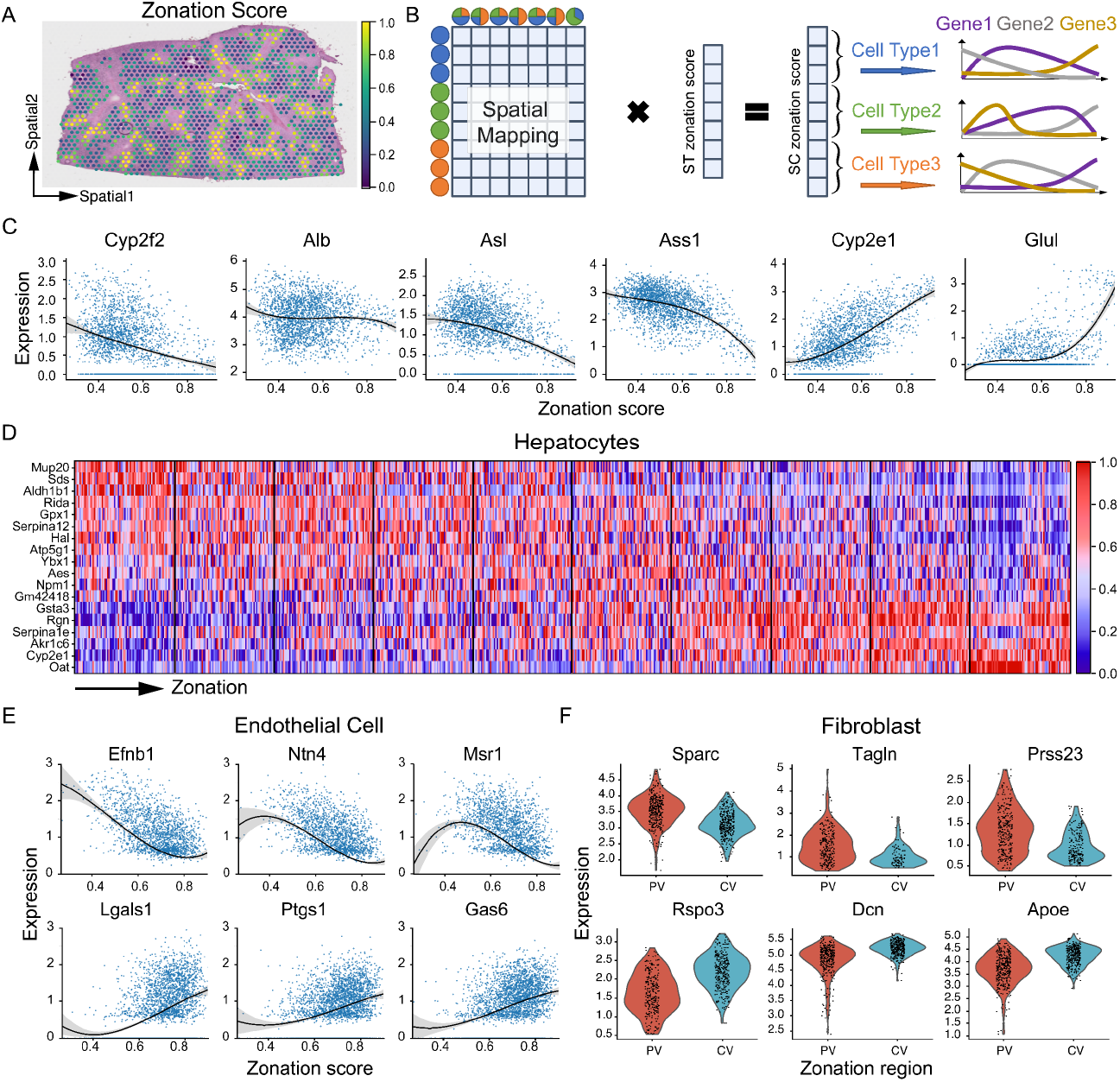
Cell-type-specific Transcriptomic Variation Along the Liver Zonation Revealed by STEM. A) Distribution of zonation scores on the ST data. A high score indicates the CV region, while a low score indicates the PV region. B) Illustration of the transfer of zonation scores from ST to SC data. The zonation scores of SC data are obtained by multiplying the SC-ST mapping matrix with the ST zonation score vector. Then cells are grouped into different cell types, and the analysis of cell-type-specific gene variation along the axis can be performed. C) Expression profiles of six zonation landmark genes along the PV-CV axis. The x-axis represents the zonation score, and the y-axis represents the gene’s raw count expression level. D) Heatmap of the top significantly differentially expressed genes along the PV-CV axis. Gene expression values are scaled, with red indicating high expression and blue indicating low expression. E) Expression profiles of six endothelial cell-specific marker genes along the PV-CV axis. The top and bottom genes are highly expressed in the PV and CV regions, respectively. F) Violin plots of fibroblast-specific marker genes identified by STEM. The top panel shows the PV marker gene, while the bottom panel shows the CV marker gene.

We first studied hepatocytes which were known to have a significant spatial variation of gene expression along the PV-CV axis^47,49^. Six reported marker genes^50^ *Cyp2f2, Alb, Asl, Ass1, Cyp2e1*, and *Glul* were highly expressed in the PV, middle, and CV region of the hepatic lobule, as shown in Fig. 5C. By further dividing hepatocytes into 10 spatial subregions along the zonation axis, 819 statistically significant regionally highly expressed genes (FDR < 0.05) were identified. For instance, Gene *Mup20, Sds* and *Aldh1b1* were highly expressed near the PV region. Gene *Ybx1, Aes* and *Npm1* were highly expressed in the middle region of the trajectory. Gene *Serpinale, Akr1c6* and *Oat* were highly expressed near the CV region. These results demonstrated STEM could potentially provide a more comprehensive set of marker genes to reveal the spatially varied transcriptomic properties of hepatocytes. We conducted a similar analysis on endothelial cells which had also been reported to have spatial variations in a recent study^48^. Two genes *Gja5* and *Adgrg6* were identified as the marker of endothelial cells at the PV region, which was in accordance with the Molecular Cartography results shown in the previous study^48^. Besides, genes such as *Lgals1, Ptgs1*, and *Gas6* were highly expressed near the PV region, and genes such as *Ntn4, Msr1*, and *Efnb1* were highly expressed near the CV region.

Then we studied the spatial variation of gene expression in fibroblasts between the PV and CV regions. Fibroblasts are critical to hepatic fibrogenesis and have attracted interest as a potential therapeutic target^51^. The transcriptomic variation among different functional sub-cell types of fibroblasts has been studied in the original study, but the variation in spatial zonation has not been investigated. Fibroblasts were categorized into three groups based on their zonation score: those near the PV region, those in the middle, and those near the CV region. The differential analysis is performed between cells near PV and CV regions, and 126 and 105 genes significantly highly expressed (FDR < 0.05) of the two regions were found, respectively. Among them, *Sparc, Tagln* and *Press23* were the most significantly highly expressed genes near the PV region. *Rspo3, DCN* and *Apoe* were the most significantly highly expressed genes near the CV region. These genes found by STEM could further help to guide gene selection for in-situ sequencing experiments. All the above results demonstrated that STEM enables assigning cells with spatial information for studying the transcriptomic variation of interested cell types along the anatomic or functional axis of tissues.

## Discussion

Revealing the spatial variation of gene expression in tissues and the spatial heterogeneity of cellular transcriptional signatures at the single-cell level is vital to understanding the functional organization of tissues and the underlying mechanisms of various diseases. Due to current limitations in sequencing techniques, computational methods for constructing spatial gene expression landscapes at the single-cell level is in need. The key is to infer the SC-SC spatial adjacency and SC-ST mapping by learning gene-spatial relations from the ST data. We proposed the method STEM for this purpose. It learns spatially-aware embeddings of transcriptomic data via transfer learning. The learned embeddings support inferring spatial adjacency between spots in ST data and cells in SC data, as well as between cells in SC data.

STEM offers unique advantages for the integrated analysis of SC and ST data. Compared to traditional spot deconvolution algorithms, STEM overcomes the limitations of fixed selections of cell-type numbers and allows for the description of cells’ continuous status or exploration of spatial distribution at different levels. Compared to other spatial mapping algorithms, our semi-simulation and biological verification experiments have shown that STEM provides more accurate spatial reconstructions that are consistent with referenced spatial topology. This enables joint analysis of the location of single cells with referenced tissue images, such as colocalizing TSK cells with the tumor leading edge. STEM also supports the analysis of transcriptomic variation within a specific cell type along the spatial axis. Using the single-cell level spatial landscape of liver tissue reconstructed by STEM, we identified gene expression changes in hepatocytes, endothelial cells, and fibroblasts. Upon the model, by following the spatial adjacency-embedding-gene pathfinding, we highlight the spatially dominant genes (SDGs) in the STEM model. These SDGs can be used for model interpretation and facilitate the discovery of healthy or diseased tissue organization mechanisms. Furthermore, Identifying SDGs of cancer cells or disease-related cell types could also be used to provide insights of potential drug targets.

STEM can receive input data of various conditions. The analysis on the MTG and hSCC datasets showed the ability of STEM to integrate SC data with different types (in-situ or sequenced based) of ST data. The mouse liver dataset demonstrated that STEM can produce convincing results when ST and SC data are not from the same donor but are from the same healthy region with similar cell type proportions. In the future, we will try to apply STEM to SC and ST data from different donors in the same disease state.

Spatial transcriptomics has become a valuable technique in investigating the biological process in different tissues, and has generated a wealth of data^52^. In the future, it is anticipated that more tissues will be characterized using both SC and ST data. Integrating these two data can provide a comprehensive understanding of cell interactions and spatial niches at single-cell resolution. We expect that STEM will be a valuable method for reconstructing single-cell spatial transcriptomic landscapes and enhancing the understanding of spatial cellular heterogeneity.

## Methods

### Data preprocessing process of STEM

STEM takes spatial gene and single-cell expression as input, and predicts the ST-ST spatial adjacency matrix at the training stage. The corresponding ground truth spatial adjacency matrix is converted from spatial coordinates. As for the input, the spatial gene expression data *X*^*ST*^ is a *N × N* matrix, where *N* is the number of spots, *H* is the number of highly variable genes identified from the standard SCANPY workflow. The single-cell gene expression data *X*^*SC*^ is a *M × N* matrix, where *M* is the number of cells. The gene is aligned with ST data to make the unified input for STEM. The gene expression value in these matrices is normalized: The unique molecular identifier (UMI) counts for each gene is divided by the total counts across all genes, and then multiplied by 10,000 and transformed into a log scale.

*Ground truth spatial adjacency matrix* Giving the ground truth ST data spatial coordinates *Y*^*ST*^ ∈ ℝ^*N*×*2*^, we convert the absolute coordinate values into a normalized spatial adjacency matrix *S*. Each element *S*_*i*_ in the matrix represents pairwise spatial association strength and is computed by the Gaussian kernel between coordinate 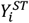 of spot *i* and 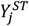 of spot *j*. The specific form of Gaussian kernel function is:

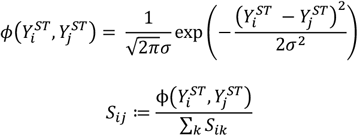

where σ is the standard deviation which controls the width of the Gaussian bell, 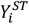 and 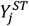 are the coordinates of two ST spots. Then L1 normalization is applied over columns in *S* to let the sum of values in each row be 1, and thus each row is the normalized distance from spot *i* to all spots. To select a proper size of σ, on the mouse embryo semi-simulation data we traversed the values of σ from 0 to 3 times of distance between two adjacent spots, and use the hit number as the evaluation metric. We found that by setting σ as the half of the adjacent distance our model achieved the highest performance.

### Encoder of STEM

STEM uses a shared MLP encoder to represent SC and ST gene expression vectors as embeddings in a unified latent space. As a spot in ST data contains more cells than a single cell in SC data, the number of expressed genes in one spot is generally higher than in a single cell, resulting in a higher sparsity of the SC data. To account for this difference in sparsity, we add a dropout layer for ST data before the encoder. The hyperparameter *d* in the dropout layer which represents the probability of an element to be zeroed is defined as:

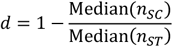

where *n*_*SC*_ and *n*_*ST*_ represent the average number of expressed genes in SC and ST data, respectively. Intuitively this dropout layer will set the sparsity of two data in the same level. Then we use an 3 layer MLP encoder to embed the expression vectors into latent embeddings Z^ST^ ∈ ℝ^h^ and *Z*^*SC*^ ∈ ℝ^*h*^ with the same dimension size *h* = 128.

### Predictor of STEM

Based on the embeddings obtained from the encoder, STEM reconstructs the spatial adjacency relationship of ST data in two ways, corresponding to the two predictor parts in the model. The first predictor utilizes only ST embeddings to reconstruct spatial relationships, while the second predictor utilizes both SC and ST embeddings.

#### Predictor part 1: spatial information extracting module

The spatial information extracting module uses ST embeddings *Z*^*ST*^ to construct the predicted ST spatial adjacency matrix 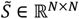:

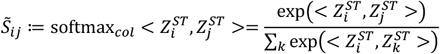

where <·,·> represents the inner products, *softmax*_*col*_ denotes the softmaxing operation over columns.

Then STEM uses the cross entropy *H* as the loss function between the ground truth and predicted ST spatial adjacency matrix 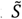 and *S*. Specifically, the row vectors 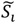 and *S*_*i*_ represents the predicted and ground truth normalized distance from spot *i* to all spots, respectively. The cross entropy loss is applied to these row vectors and can be described as:

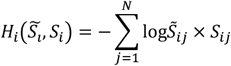

The total cross entropy loss is defined as the mean of all rows’ loss:

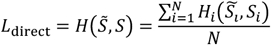

#### Predictor part 2: domain alignment module

In this part, STEM first reduces the mean distance between ST and SC embeddings, and then uses these embeddings to estimate the SC-ST and ST-SC mapping matrices. These two mapping matrices are multiplied together to construct another ST spatial adjacency matrix 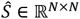 By directly optimizing 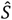, the optimal SC-ST and ST-SC mapping relationship can be found. A similar idea is proposed in Haeusser’s work^53,54^. We extend its applicability from classification to relation construction and fully utilize the cross-domain association matrix as the SC-ST and ST-SC mapping matrix.

Specifically, the STEM introduces the Maximum Mean Discrepancy loss to reduce the mean distance of ST and SC embeddings:

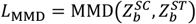

where 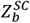 and 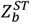are SC and ST embeddings in a mini-batch.

The inner-product is used to measure the similarity between ST and SC embeddings: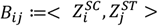. The mapping matrix C ∈ ℝ^M×N^ from SC to ST is computed by softmaxing similarity matrix *B* over columns:

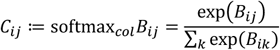

Similarly, the mapping matrix from ST to SC 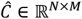 is computed in the same way by replacing *B* with *B*^T^. Then the two-step spatial adjacency matrix 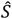 is the multiply results of two mapping matrices:

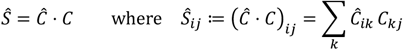

The cross entropy *H* between the ground truth and this two-step ST spatial adjacency matrix is used as the loss function:

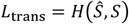

The total loss of STEM consists of three parts:

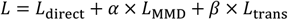

We set the hyperparameter α= *0*.5 and gradually increase *β* from 0 to 1 during the training stage. The parameters in the STEM model are optimized by minimizing total loss *L* using stochastic gradient descent with momentum. The increasing weight *β* encourages STEM first to focus on reconstructing the spatial adjacency within ST and then learn the spatial mapping between ST and SC.

After training, STEM gives multiple results including the cell type deconvolution results *T*^*ST*^∈ ℝ^N×D^, pseudo spatial coordinates of single cells *Y*^*SC*^∈ ℝ^M×*2*^ and the SC spatial adjacency matrix *S*^*SC*^ ∈ ℝ^M×M^. STEM uses the ST-SC mapping matrix *Ĉ* for deconvoluting the cell type proportion in each spot. Given a cell type indicator matrix *T*^*SC*^∈ ℝ^N×D^, where D is the number of cell types. Each row in *T* is a one-hot vector, and the nonzero index is the corresponding cell type. The deconvolution result *T*^*ST*^ ∈ ℝ^M×D^ is given as:

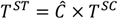

The pseudo spatial coordinates of single cells are retrieved by computing:

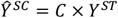

And the spatial adjacency matrix is the inner-product results of SC embeddings:

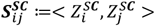

### Attribution function

STEM employs the integrated gradient (IG) technique for each cell to assign attribution of genes to its desired spatial location. The IG technique is based on counterfactual intuition, which considers the absence of the cause as a baseline and compares the baseline with current results. For a computational model, the baseline absence of the cause is modeled as a zero input vector. Specifically in STEM, let *X*_*i*_ be the input gene expression of cell *i, C*_*i,m*_ is the max value in matrix *C*’s *i*th column, and *X*^*′*^ = [*0,0*, …, *0*] is the baseline vector. Given this three information, the attribution *W*_*ij*_ of gene *j* in cell *i* is computed as the integrated gradient along the path from the baseline *X*^*′*^ to the input *X*_*i*_:

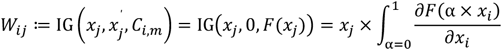

As the maximum value *C*_*i,m*_ indicates that cell *i* has the maximum probability located around ST spot *m*, the attribution vector *W*_*i*_ reveals the contribution of genes for determining this spatial location. We compute the gene attribution vector of all single cells and get a gene attribution profile *A*^M×*H*^.

### Semi-simulation data generation

Transcriptomic data with spatial information at the single-cell resolution is required for evaluating the methods’ ability of inferring spatial associations. Currently, such data can only be provided by some less popular spatial sequencing technologies which are low-throughput and require complicated operations. We take these experimental single-cell resolution spatial transcriptomic data as SC data with the ground-truth spatial information, and simulate pseudo-ST data by creating a spatial grid on the spatial space of these data.

Specifically, we placed the pseudo-ST spots at the crossing point of the spatial grid. The number of pseudo-ST spots are decided by the topology structure and spatial distribution of the original data. For example. an 30 × 40 grid is generated for mouse embryo data. The gene expression profile of each spot is obtained by summing the expression of its surrounding single cells. We kept the spots which contains more than 3 cells. It is noticeable that in real spatial data the tissue cannot be fully covered by spots, so the transcripts of some single cells cannot be captured. We take this into account in the simulation process. The gene expression of each spot aggregates only about 50-70% of the local surrounding single cells. In other words, one-third of single cells gene expression profiles are not included in the ST data. Then the cell type proportion of each spot is calculated based on its contained single cells annotation. Pseudo-spots in the simulated ST data have the information of gene expression *X*^*ST*^, spatial coordinate *Y*^*ST*^ and cell type proportion *P*, where *P* is a *N* × *A* matrix and *A* is the number of cell types appeared in the tissue.

### Output unification

To make the results comparable, we unify the outputs of different methods into three parts: the SC spatial adjacency matrix, the reconstructed SC spatial coordinates and the SC-ST mapping matrix. For methods that predict SC coordinates (CellTrek and Seurat), the SC-ST mapping matrix and SC adjacency matrix are obtained by calculating the spatial distance between or within single cells and spots. The closer the spatial distance between the cells or spots, the higher the weight in the matrix. For other methods that provide the SC-ST mapping matrix (Tangram and Spaotsc), the SC coordinates are obtained by averaging the coordinates of spots according to the mapping weights.

### Performance evaluation

We validate the model performance by using three metrics on the synthetic data. We compute the mean absolute error (MAE) of distance between the predicted and true spatial coordinates as the error metric. We used the hit number to justify the correctness of predicted SC-SC adjacency. We calculated the Pearson correlation coefficient (PCC) between the predicted and true cell type spatial distribution.

*MAE* The predicted coordinates are computed by multiply the SC-ST mapping matrix with ST spatial coordinate vector. Distance MAE is defined as the mean distance among all predicted and ground truth coordinate pairs:

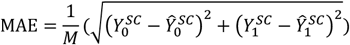

where *Y*_0_^·^ and *Y*_1_^·^ are the coordinate values in the first and second spatial axis.

*Hit number* Hit number is calculated on the SC-SC spatial adjacency matrix. It measures the average of number of cell’s K-nearest neighbors that can be successfully predicted, and it is the function of K. the hit number is increased the max hit number is equal to K when K is the number of all cells:

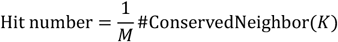

*PCC* For cell type *i*, it has a spatial proportion distribution among all ST spots. The distribution can be flattened into one column as *P*_*i*_. The PCC is computed between the estimated cell distribution vectors 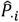 and ground truth:

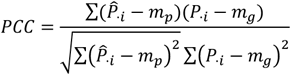

where *m*_*p*_ is the mean of the vector 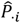 and *m*_*g*_ is the mean of the vector *P*_*i*_.

### Data processing

#### Mouse Embryo

We obtained the raw count matrices and selected the “z=2” slice for each embryo section. We removed cells with “Low quality” annotation. We used the raw count matrices to generate the pseudo ST data. And then we normalized and log scaled the SC and generated ST data by using the “normalize_total” and “log1p” function of SCANPY package in python.

#### Human Middle Temporal Gyrus

We used the 4000-gene MERFISH data from the donor “H18” as the ST data. We manually separated the MERFISH data into L1-L6 layers according to the figure 3 shown in the original paper. We used the exons matrix of SMART-seq data as the SC data. We categorized cells into different layers according to the “brain_region” column in the provided cell metadata. After gene alignment, 3,491 genes were shared across the SC and generated ST data.

#### Human squamous cell carcinoma

We used single-cell RNA-seq and the Spatial Transcriptomic data of donor 2 and donor 10. We removed single cells with “Multiplet” annotation. To reduce the computation burden, we selected 2,000 highly variable genes from the ST data as the common gene set for SC and ST gene alignment.

#### Mouse liver

We utilized four Visium liver sections from mouse sample 1 as the ST data. We used both CD45- and CD45+ cells (sample ‘CS88’,’CS89’,’CS93’,’CS97’,’CS138’ and ‘CS141’) sequenced by scRNA-seq as the SC data. As the number of Endothelial cells and Kupffer cells in the SC data was three times higher than other cell types, we implemented a subsampling strategy with a sampling rate of 0.3. We then identified 2,000 highly variable genes in each Visium section and merged them into a union common gene set containing a total of 4866 genes. We used this gene set to align the genes between the SC and ST data.

## Data availability

The original data used in this paper can be accessed through the following links. The mouse embryo data can be downloaded from https://marionilab.cruk.cam.ac.uk/SpatialMouseAtlas. The human MTG MERFISH data can be downloaded from https://datadryad.org/stash/dataset/doi:10.5061/dryad.x3ffbg7mw, and the SMART-seq data can be downloaded from https://portal.brain-map.org/atlases-and-data/rnaseq/human-mtg-smart-seq. The hSCC ST and SC data can be obtained from the GEO database (GSE144240). The mouse liver ST and SC gene expression data can be download from the “Liver Cell Atlas: Mouse StSt” dataset in https://www.livercellatlas.org/download.php. The corresponding spatial coordinates and corresponding H&E images can be obtained from the GEO database (GSE192742).

## Code availability

An open-source implementation of the STEM algorithm can be downloaded from https://github.com/WhirlFirst/STEM.

## Acknowledgements

This work was partially supported by National Natural Science Foundation of China (NSFC) (grants 62250005, 61721003 and 62103227), National Key R&D Program of China (grant 2021YFF1200901) and Tsinghua-Fuzhou Institute for Data Technology (TFIDT2021005).

## Author contributions

M.H., L.W., and X.Z. conceived the study. M.H. E.L. and Y.W. collected datasets involved in this article. M.H. and E.L. benchmarked all methods. M.H. designed and implemented the STEM algorithm. Y.C., C.L., S.C., H.G., H.B. and L.W. provided a lot of advice on algorithm implementation and simulation experiments. M.H. L.W. and Y.W. designed the biological applications. M.H., E.L., L.W. and X.Z. wrote the manuscript. All authors read and approved the final manuscript.

## Additional information

Supplementary Figures 1-11.

