## Supplementary Figures 1-11 for "STEM: A Method for Mapping Single-cell and Spatial Transcriptomics Data with Transfer Learning"

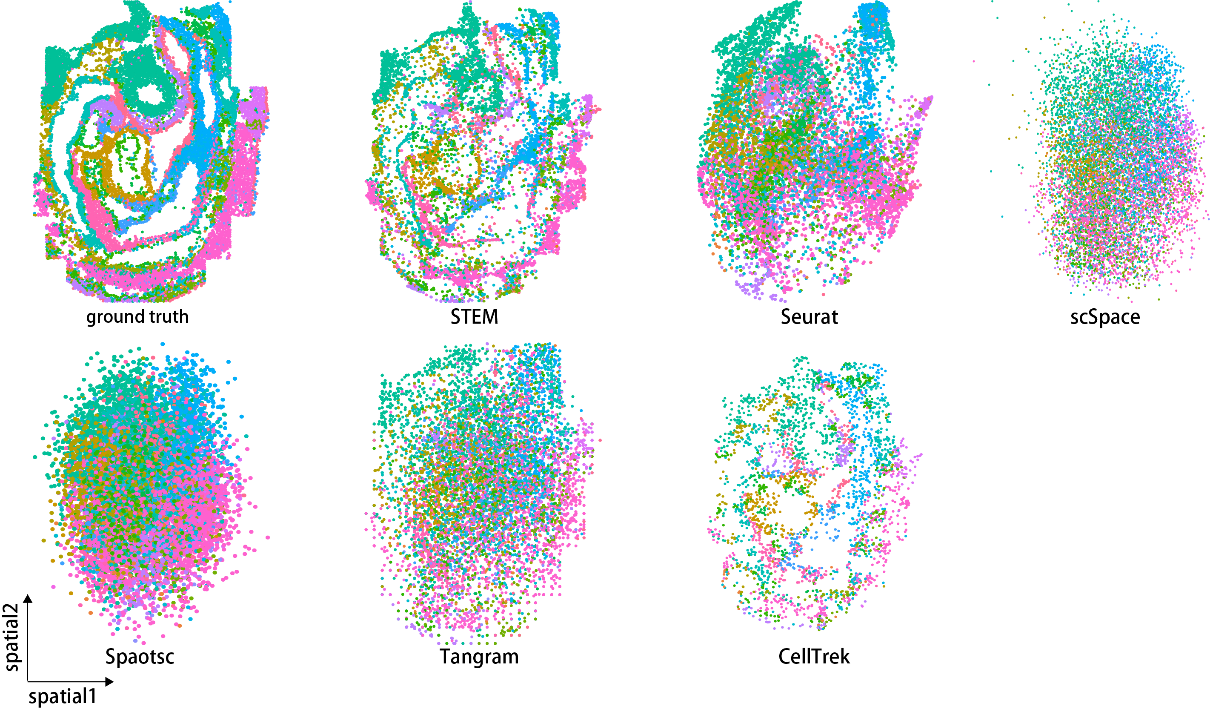

Figure S1 the spatial reconstruction results of all methods on mouse embryo 1.

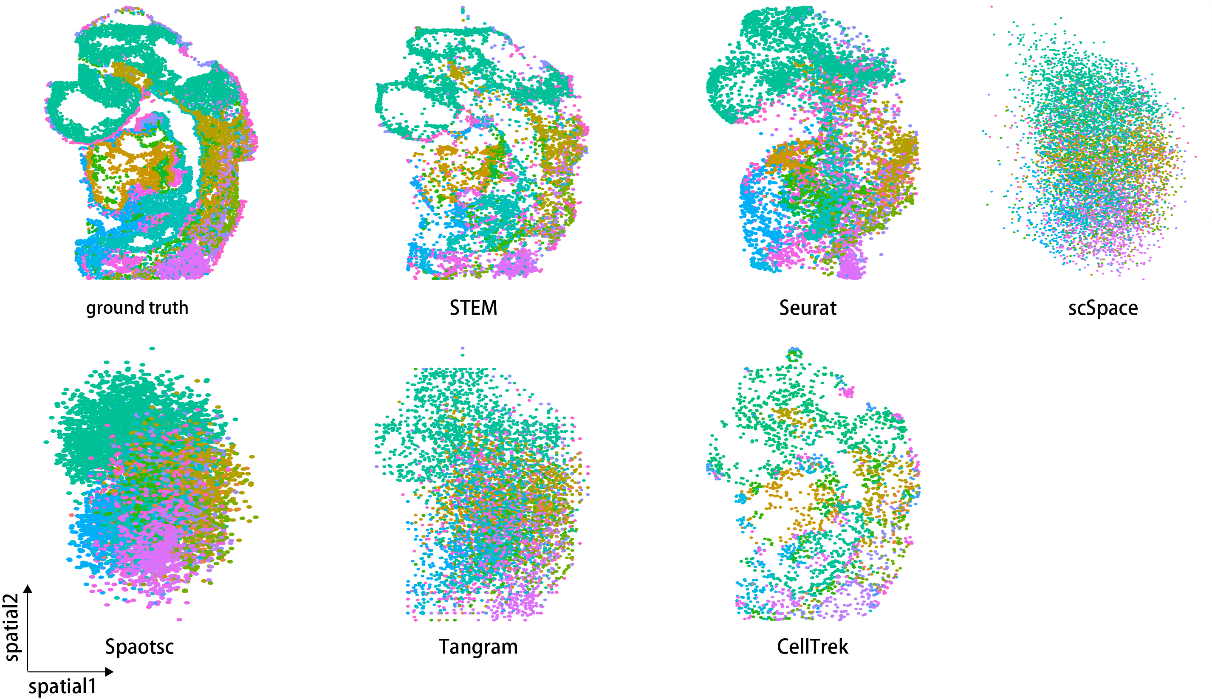

Figure S2 the spatial reconstruction results of all methods on mouse embryo 2.

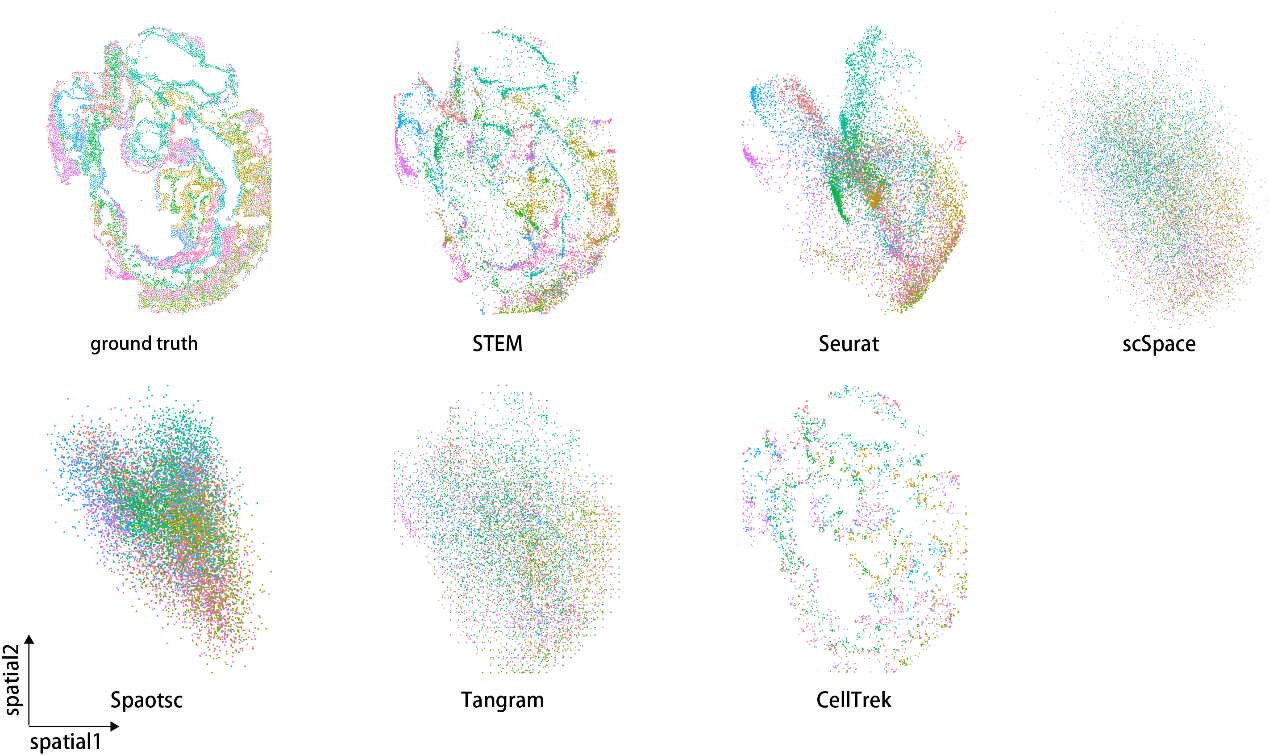

Figure S3 the spatial reconstruction results of all methods on mouse embryo 3.

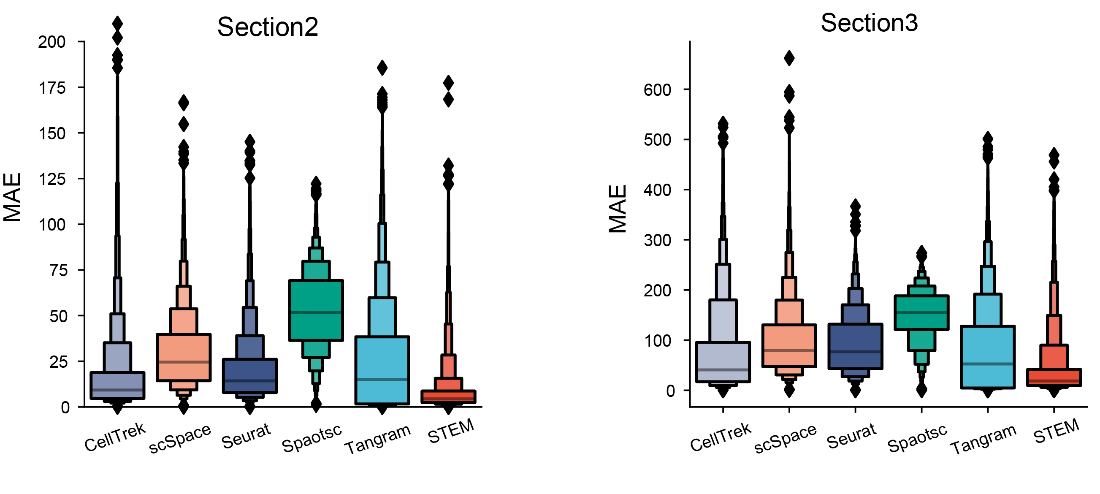

Figure S4 The means absolute error (MAE) computed by different methods’ results on the mouse embryos 2 and 3.

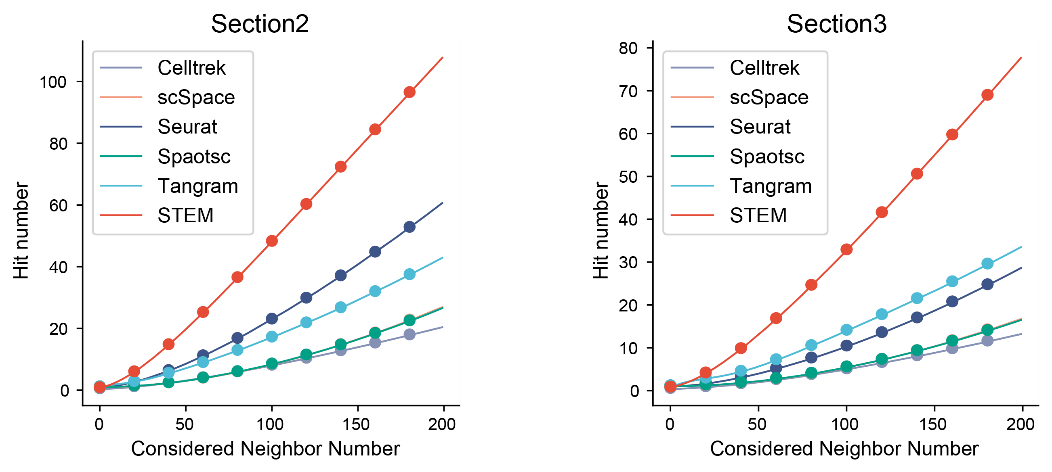

Figure S5 The Hit number computed by different methods’ results under different consider neighbor numbers on the mouse embryos 2 and 3.

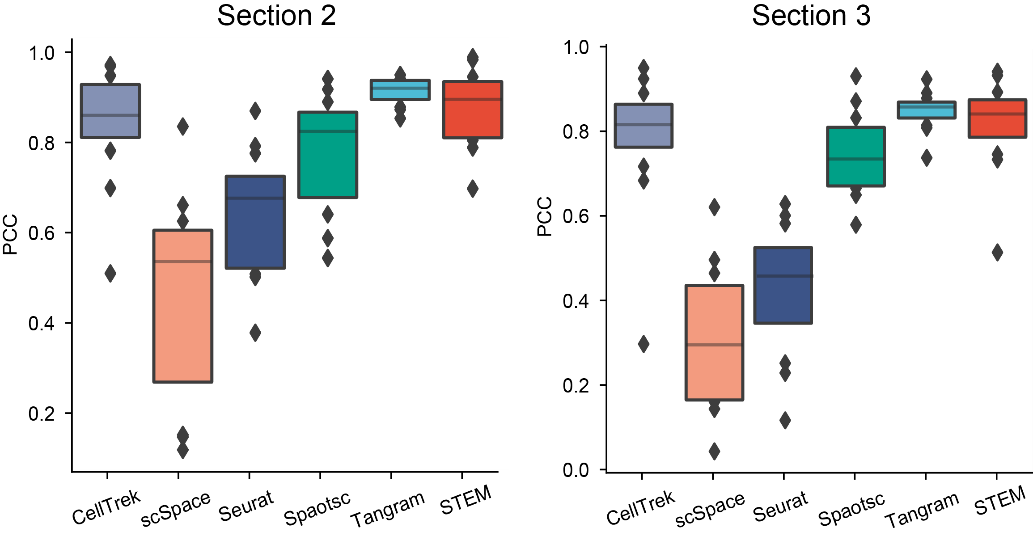

Figure S6 The Pearson correlation coefficient (PCC) computed by different methods’ results on the mouse embryos 2 and 3.

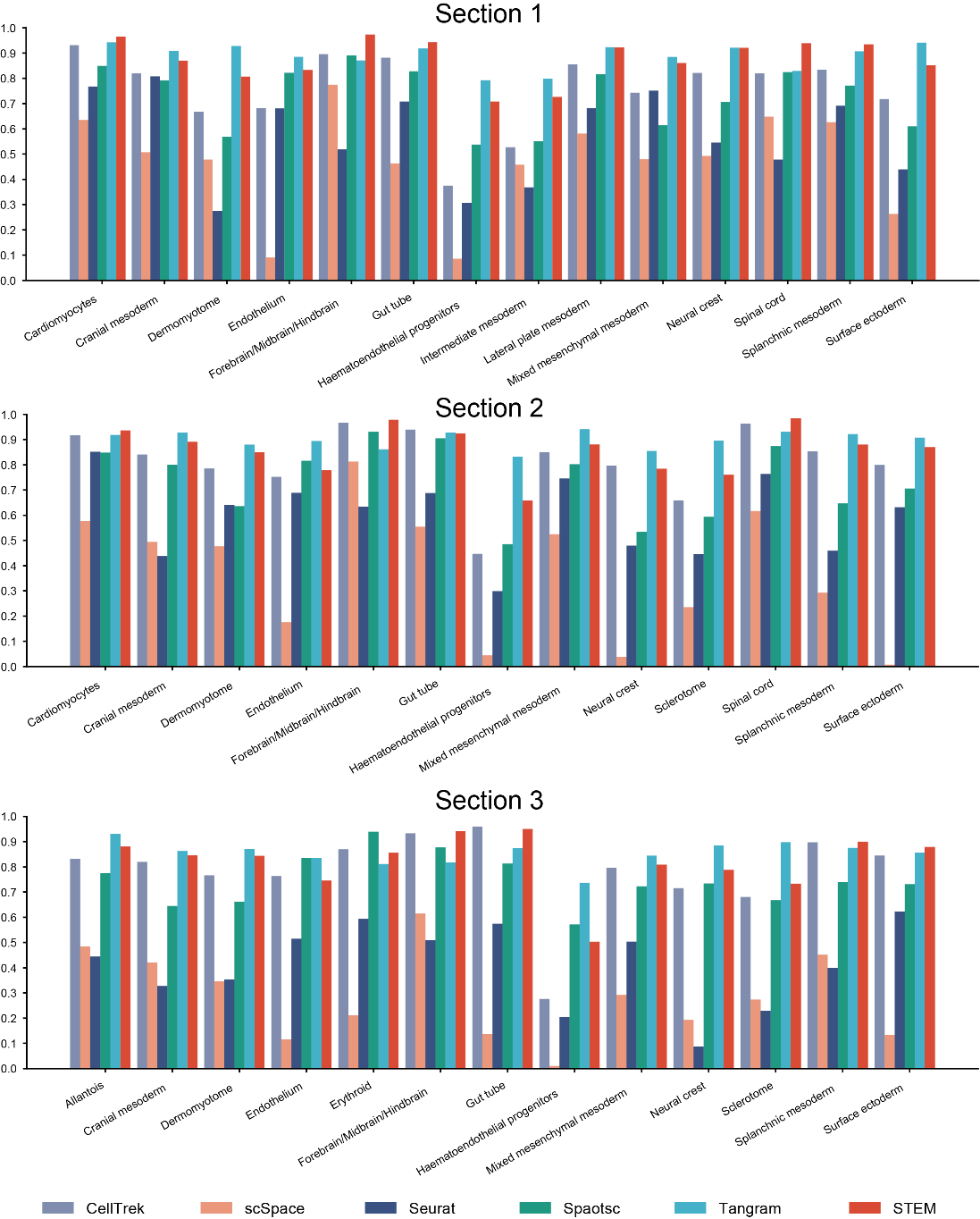

Figure S7 The PCC computed by different methods’ results among different cell types on the mouse embryo 2 and 3 data.

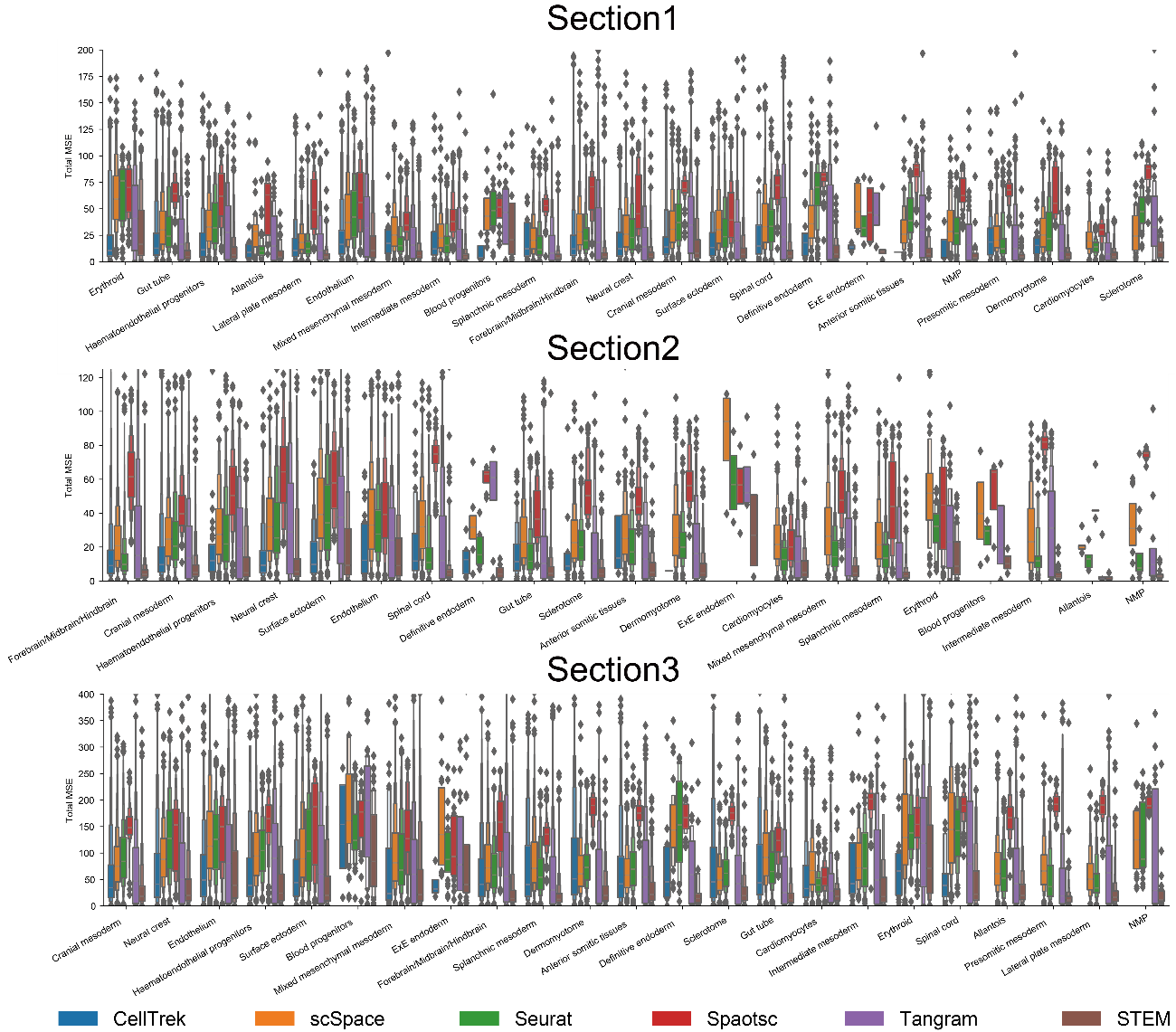

Figure S8 The MAE computed by different methods’ results among different cell types on all mouse embryo data.

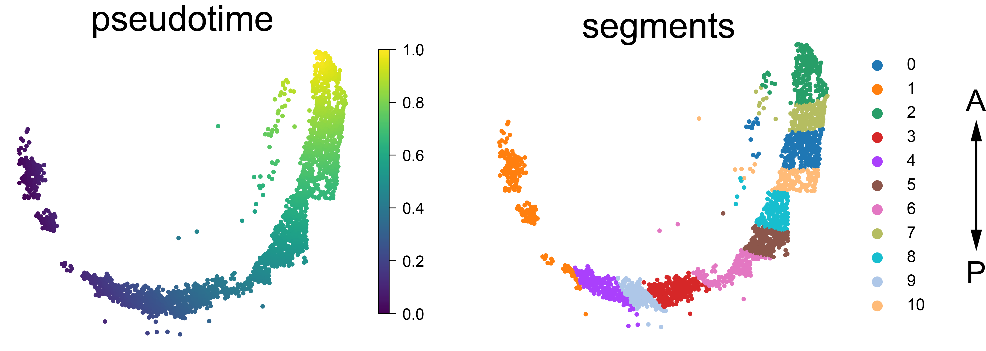

Figure S9 The pseudo-time trajectory and segments generated in the spinal cord region. A: anterior, P: posterior

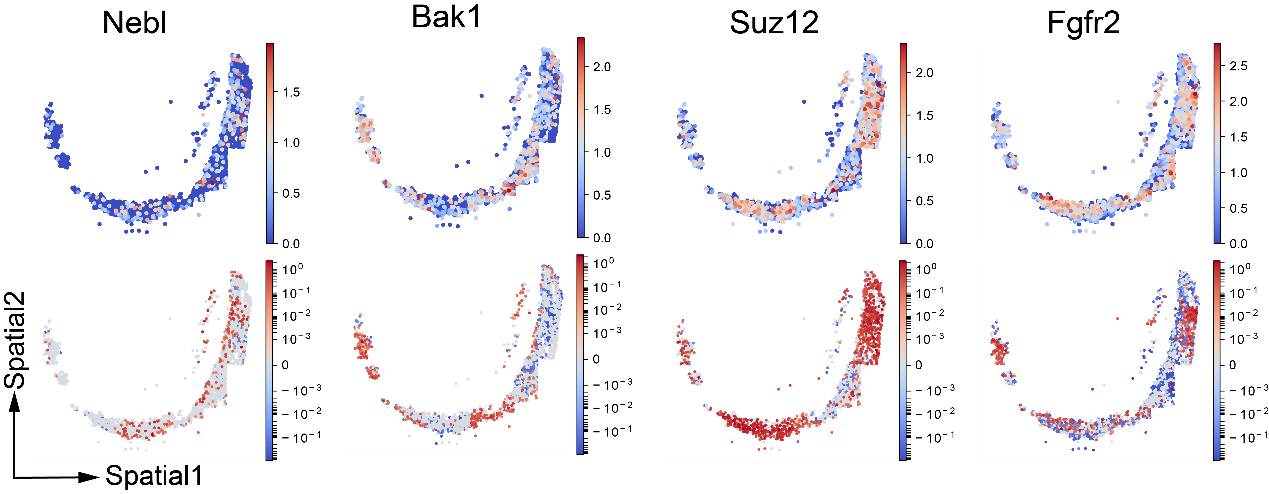

Figure S10 Gene expression and attribution score spatial pattern in spinal cord region. The attribution score is shown in the log form for better visualization. Above: Gene expression. Bottom: Gene attribution score

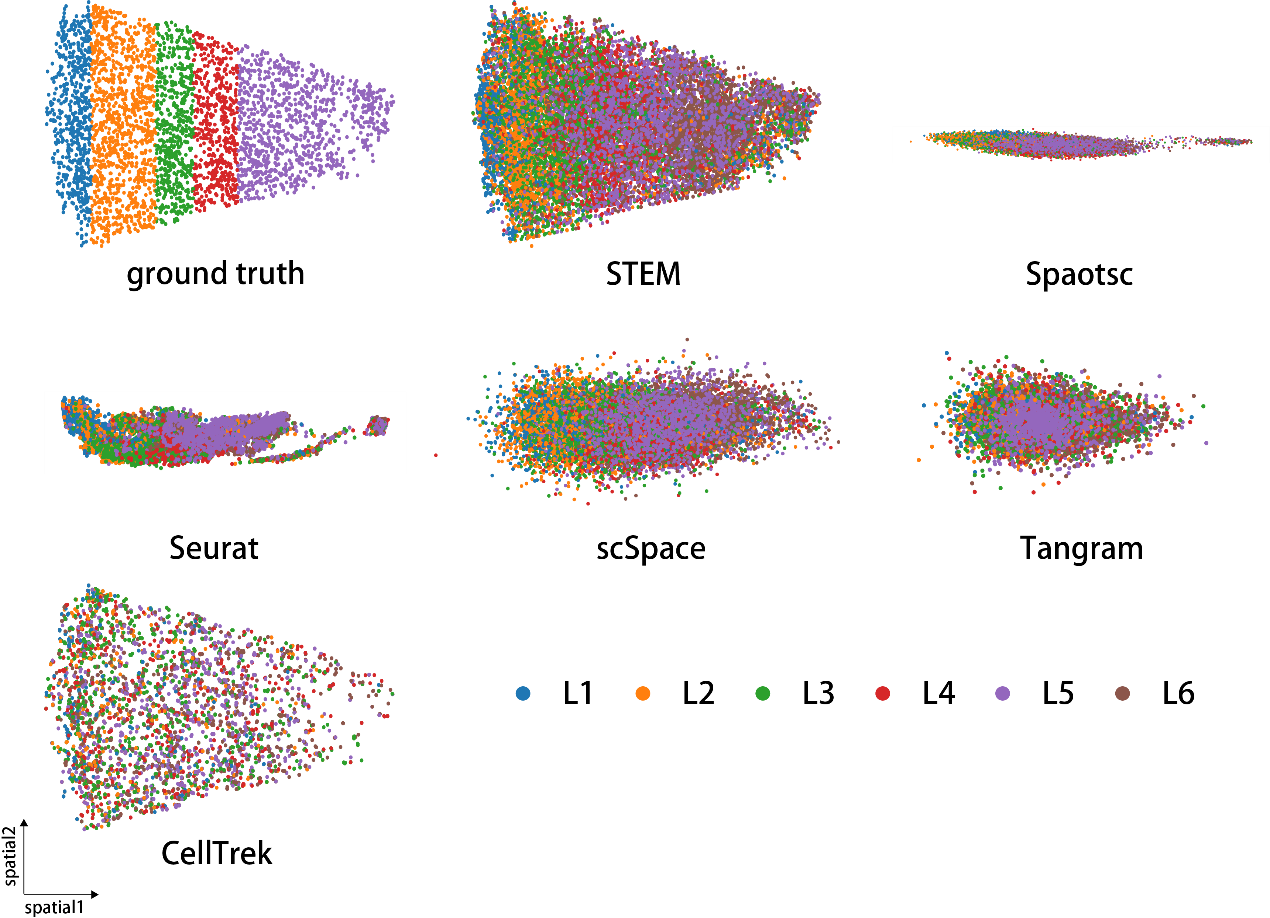

Figure S11 the spatial reconstruction results of all methods on human MTG data.
